# The lncRNA *Gm16685*/*MITA1* modulates inflammatory astrocyte reactivity through PCBP2 associated regulation of IKKβ signaling

**DOI:** 10.64898/2026.07.03.736437

**Authors:** Ulrike Fuchs, Sophie Schröder, Tonatiuh Pena, Dennis M Krüger, Susanne Burkhardt, Anna-Lena Schütz, Farahnaz Sananbenesi, André Fischer

## Abstract

Long non-coding RNAs (lncRNAs) are increasingly recognized as regulators of cellular identity and disease associated gene expression programs, yet their role in astrocyte reactivity remains poorly understood. Here, we profiled lncRNA expression in primary mouse astrocytes exposed to inflammatory activation paradigms that model microglia driven signaling. This identified a conserved set of activation responsive lncRNAs, among which *Gm16685* emerged as one of the most strongly induced candidates. *Gm16685* and its human homolog *MITA1* were enriched in the nucleus, and *MITA1* expression was increased in selected human datasets from Alzheimer’s disease, Parkinson’s disease and frontotemporal dementia patients. Functional depletion of *Gm16685* attenuated inflammatory gene expression and several activation associated astrocyte phenotypes, including reactive oxygen species production, glutamate handling, phagocytic activity and proliferation. Time-resolved transcriptomic analysis indicated that *Gm16685* is required for the timely induction of inflammatory response genes. Mechanistically, *Gm16685*/*MITA1* interacted with the RNA binding protein PCBP2, and *Gm16685* depletion was associated with reduced PCBP2 protein abundance, altered splicing of Inhibitor of NF-κB Kinase Subunit Beta (IKKβ) and a shift in downstream inflammatory signaling. Together, our findings identify *Gm16685*/*MITA1* as a conserved lncRNA regulator of astrocyte reactivity and suggest that non-coding RNA dependent control of RNA binding proteins contributes to inflammatory signaling in neurodegenerative disease relevant contexts.

## Introduction

Long non-coding RNAs (lncRNAs), defined by a length greater than 500 nucleotides and a lack of protein-coding potential (Mattick et al., 2023), have emerged as central regulators of gene expression in the central nervous system (CNS), where they fine-tune transcriptional and post-transcriptional programs across development, homeostasis, and disease (Altaf et al., 2025). Consequently, altered lncRNA expression profiles have been associated with a broad spectrum of neurological and neurodegenerative diseases (Esmaeili et al., 2025), positioning lncRNAs as key components of pathogenic gene-regulatory networks and potential therapeutic entry points.

In parallel, there is increasing recognition that glial cells are active participants in the initiation and progression of neurodegeneration. Astrocytes constitute the most abundant type of glial cells in the CNS that continuously survey the neural environment and respond to pathogens or other injury signals by adopting reactive states that profoundly reshape their transcriptional, metabolic and secretory profiles (Li et al., 2019), (Patani et al., 2023), (Lawrence et al., 2023). While transient glial activation can be protective, chronic reactive states contribute to synaptic dysfunction, neuronal excitotoxicity, and blood-brain barrier leakage, amplifying neuronal vulnerability in a feed-forward manner (Liddelow et al., 2017).

Recent studies have begun to uncover that lncRNAs are integral components of the regulatory circuitry governing astrocyte reactivity and their contribution to neurodegenerative diseases. Astrocyte-enriched lncRNAs control transcriptional programs linked to neuronal support, synapse organization and inflammatory signaling, with specific examples becoming dysregulated in aging and Alzheimer’s disease (AD) models, modulating reactivity or cognitive decline (Chen et al., 2021), (Schroeder et al., 2024), (Schröder et al., 2024). However, the identities and functions of many lncRNAs induced during astrocyte activation remain poorly characterized.

To address this gap, we performed a comprehensive transcriptomic analysis of deregulated lncRNAs in primary mouse astrocytes following activation with inflammatory stimuli that model key features of the neurodegenerative microenvironment (Liddelow et al., 2017), namely lipopolysaccharide (LPS)-stimulated microglia conditioned medium (MCM) or the cytokine mix of Interleukin 1 alpha (IL-1α), tumor necrosis factor alpha (TNF-α) and complement component 1q (C1q) (often referred to as TIC) that mimics this response. Both conditions elicit a highly overlapping response regarding protein-coding genes and lncRNAs, and we hypothesized that lncRNA perturbations would affect reactive astrocyte phenotypes.

The lncRNA *Gm16685* emerged as a prime target due to its robust upregulation and evolutionary conservation as well as its described role in Nuclear Factor kappa Beta (NF-κB)-mediated inflammatory responses in other tissues (Akıncılar et al., 2021). We investigated its loss during activation in mouse astrocytes and found that *Gm16685* knockdown attenuated core inflammatory responses. We identify *Gm16685*/*MITA1* as a conserved, activation induced lncRNA that modulates inflammatory astrocyte phenotypes and provide evidence linking it to PCBP2 dependent regulation of *Ikbkb* mRNA processing and downstream inflammatory signalling. Taken together, this study elucidates non-coding mechanisms in glial cells and identifies reactive astrocyte expressed lncRNAs as candidate regulators of inflammatory signalling in disease relevant contexts.

## Results

Our aim was to systematically investigate the role of lncRNAs in microglia-driven astrocyte activation. Therefore, primary mouse astrocytes were treated for 24 hours with either conditioned medium from 6-hour LPS-activated microglia or with the TIC cytokine mix, followed by total RNA sequencing (**Fig. 1A**, **Fig. S1A, Supplementary tables 1 and 2**). Both treatments elicited highly similar transcriptional responses, with approximately 70 % overlap in differentially expressed genes (60% for lncRNAs) (**Fig. 1B**, **S1B**), and gene ontology enrichment analysis revealed an association with defence and innate immune response processes (**Fig. S1C, Supplementary table 3**).

**Figure 1.**
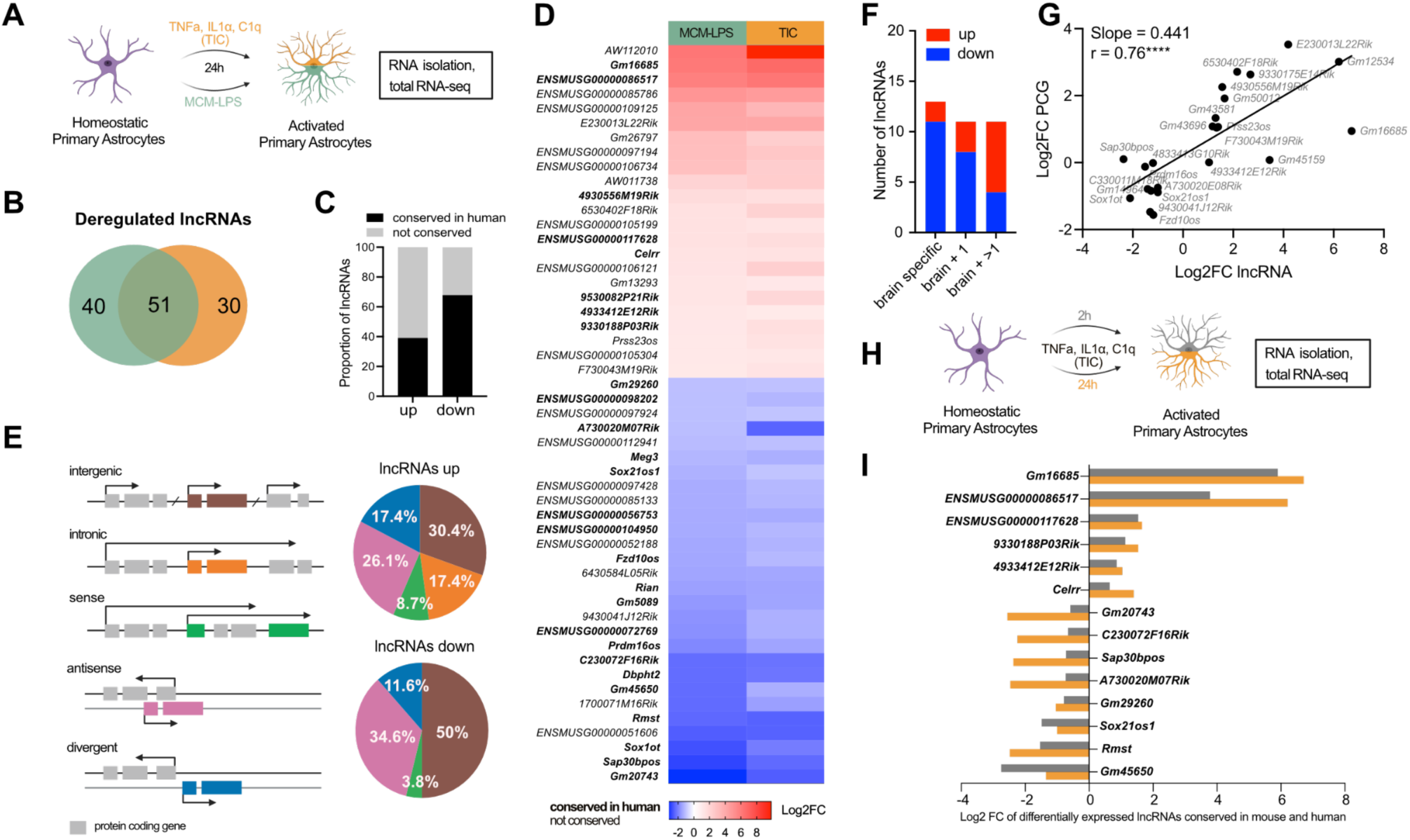
Characterization of long non-coding RNAs deregulated in activated astrocytes. **A** Schematic workflow, homeostatic mouse primary astrocytes were activated for 24 hours with microglia-conditioned medium (MCM) from 6 hour lipopolysaccharide (LPS; MCM-LPS) activated mouse primary microglia or the cytokine mix of TNF-ɑ, IL-1ɑ and C1q (TIC) followed by total RNA sequencing. **B** Venn diagram showing overlap between long non-coding RNAs differentially expressed in MCM-LPS or TIC vs. vehicle identified from total RNA sequencing (adjusted p-value < 0.05, |log2FC| > 1 and BaseMean > 25). **C** Bar plot showing proportion of activated-astrocyte deregulated lncRNAs with evolutionary conservation identified by synteny in rodents and humans identified in (B). **D** Heatmap depicting log2 fold changes (Log2FC) of lncRNAs identified in (B), conserved lncRNAs in humans and rodents are depicted in bold. **E** Schematic illustration of classification by genomic localisation in regard to DNA strand and protein coding genes. Right panel: Pie chart depicting classification of lncRNAs deregulated in activated primary astrocytes. **F** Expression of activated-astrocyte deregulated lncRNAs across different mouse organs based on data from the EMBL EBI expression atlas (https://www.ebi.ac.uk/gxa/genes/). Bar plot depicting number of lncRNAs showing a brain specific, brain and one other tissue (+1) or brain and multiple other tissues (+ >1) expression pattern. **G** Scatter plot of log2 fold change (FC) for n = 23 paired lncRNA-protein coding genes (PCG) (vicinity <100 kb) from total RNA sequencing data of TIC activated astrocytes. Linear regression slope=0.441 indicated by line (SE = 0.087, *P* = 0.0001, R² = 0.55); Spearman ρ = 0.76 (\*\*\*\**P* < 0.0001), positive correlation supports cis-regulatory role. **I** Bar plot of conserved lncRNA Log2 fold changes (FC) after 2 hours (early signalling response) vs. 24 hours (sustained reactive state) TIC activation in primary astrocytes by total RNA-seq (adjusted p-value < 0.05, |log2FC| > 0.585 and BaseMean > 25).

Next, we assessed evolutionary conservation (synteny), genomic features and tissue expression patterns of deregulated lncRNAs. Downregulated lncRNAs showed higher conservation across species (19 out of 28) compared to upregulated lncRNAs (8 out of 23) (**Fig. 1C, Supplemental table 4**); ∼50% of the downregulated lncRNAs were intergenic (18%; upregulated lncRNAs), whereas intronic lncRNAs were exclusive to the upregulated subset (12%; **Fig. 1E**), and downregulated lncRNAs were more brain-enriched (11 out of 13) (**Fig. 1F**). Given that lncRNAs have been shown to regulate proximal protein-coding genes in *cis*, we examined their concordant deregulation and observed a significant positive correlation (**Fig. 1G**).

To capture immediate inflammatory signalling (Rodgers et al., 2020), we included a 2-hour TIC activation timepoint and analysed lncRNA expression by total RNA sequencing as before (**Fig. 1H, Supplemental table 5**). We observed that half (14) of our identified conserved lncRNAs also showed at least 50 % increase or decrease (log2 fold change |0.585|) at the 2-hour timepoint (**Fig. 1I**). Collectively, these findings indicate that lncRNA expression changes broadly track those of protein-coding genes during astrocyte activation, supporting the role of lncRNAs in modulating astrocytic responses to inflammatory stimuli.

### Conserved lncRNA *Gm16685*/*MITA1* is strongly induced in reactive astrocytes and neurodegenerative diseases

To further elucidate the functional roles of lncRNAs in activated astrocytes, we focused on a conserved lncRNA showing the strongest induction at 2 hour and 24-hour TIC activation, namely *Gm16685* (**Fig. 1I**). Mouse *Gm16685* and its human homolog *MITA1* are both antisense to the IL7 coding gene, sharing 41 % global sequence similarity (**Fig. 2A**). Of note, our previous analysis revealed little correlation between *Gm16685* and *IL-7* mRNA levels (**Fig. 1G**). We validated the induction and localisation of *Gm16685* in primary astrocytes and *MITA1* in human immortalized astrocytes following 24 hour TIC activation (**Fig. 2B, C**) and found that both are significantly enriched in nuclear fractions, with a further shift to nuclear localisation in activated primary astrocytes, indicating a role in transcription. Since astrocyte activation is a hallmark of many neurodegenerative diseases, we queried publicly available sources like the AGORA database (https://agora.adknowledgeportal.org/), which includes over 1000 postmortem brain samples from both control individuals and AD patients. Here, we found *MITA1* significantly upregulated in the parahippocampal gyrus and superior temporal gyrus regions (**Fig. 2D**). In addition, increased *MITA1* expression was reported in idiopathic PD patient iPSC-derived midbrain organoids from the LncExpDB database (https://ngdc.cncb.ac.cn/lncexpdb/) as well as in different subtypes of FTD compared to non-demented controls from the RiMod FTD database (https://www.rimod-ftd.org) (**Fig. 2D**). In line with this, *Gm16685* remains significantly upregulated after 7 days of continuous TIC activation (**Fig. S2A**). Taken together, these data show the broad induction of *MITA1* in neurodegenerative diseases which share the hallmark of neuroinflammation in disease development.

**Figure 2.**
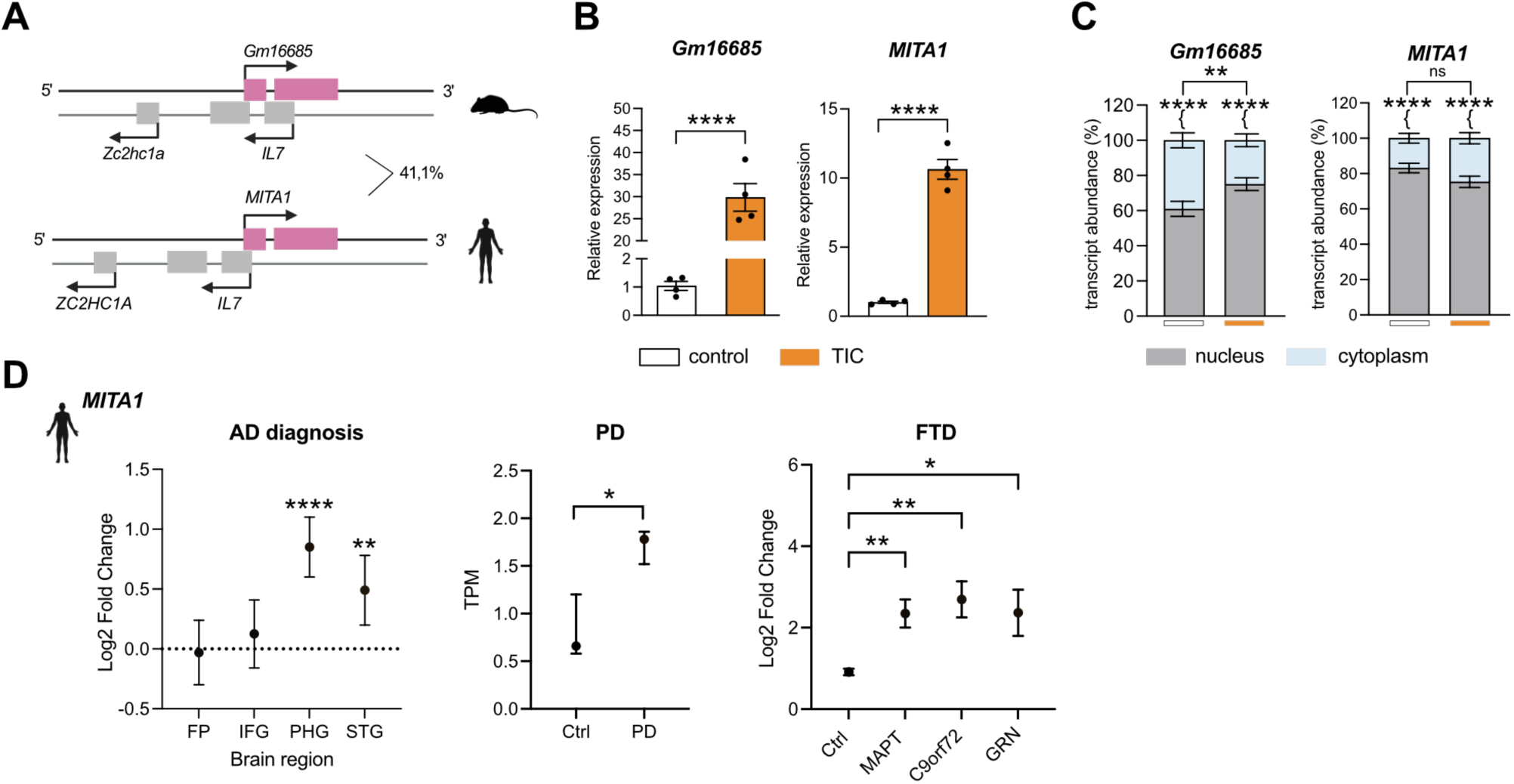
Evolutionary conserved lncRNA *Gm16685*/*MITA1* is upregulated in activated astrocytes and neurodegenerative diseases. **A** Schematic illustration showing the genomic localization of *Gm16685* (mouse chromosome 3) and its human homolog *MITA1* (human chromosome 8), classifying both as antisense lncRNA. Percentage indicates global sequence similarity. **B, C** Bar plots showing *Gm16685* expression in mouse primary astrocytes (left panel) or human *MITA1* expression in immortalized human astrocytes (right panel) activated 24 hours with the cytokine mix TNF-ɑ, IL-1ɑ, C1q (TIC) in total cells (B) or nuclear and cytoplasmic fractions (C), (one-way ANOVA; \*\*\**P* < 0.001, \*\*\*\**P* < 0.0001, ns not significant). Gene expression was normalized to 18S. **D** Dot plots showing expression changes of *MITA1* in different neurodegenerative diseases. Left panel: Log2 Fold change in different brain regions in Alzheimer’s disease (AD) patients compared to controls based on data from the Agora database (https://agora.adknowledgeportal.org/); FP frontal pole; IFG inferior frontal gyrus; PHG parahippocampal gyrus; STG superior temporal gyrus. Middle panel: Transcripts per million (TPM) in healthy controls (Ctrl) or idiopathic Parkinson’s disease (PD) patient induced pluripotent stem cell-derived midbrain organoids from the LncExpDB database (https://ngdc.cncb.ac.cn/lncexpdb/; Version 2.0). Right panel: Log2 Fold change in different subtypes of Frontotemporal dementia (FTD) patient brains compared to non-demented controls (Ctrl) based on data from the RiMod FTD database (https://www.rimod-ftd.org), (\**P* < 0.05,\*\**P* < 0.01, \*\*\*\**P* < 0.0001). Error bars represent SEM.

### *Gm16685* knockdown inhibits reactive phenotypes and delays treatment induced gene expression changes

We hypothesised that *Gm16685* loss would cause reactive astrocyte phenotype perturbations. Therefore, we performed a 48-hour antisense Gapmer-mediated knockdown (KD) of *Gm16685* in primary astrocytes including 24-hour TIC activation (**Fig. 3A**) and assessed functional phenotypes. We observed that in activated *Gm16685* KD astrocytes, glutamate uptake, phagocytic activity and proliferation resembled levels of non-activated astrocytes, while reactive oxygen species (ROS) levels were even further reduced in activated *Gm16685* KD astrocytes compared to homeostatic levels (**Fig. 3C-F**).

**Figure 3.**
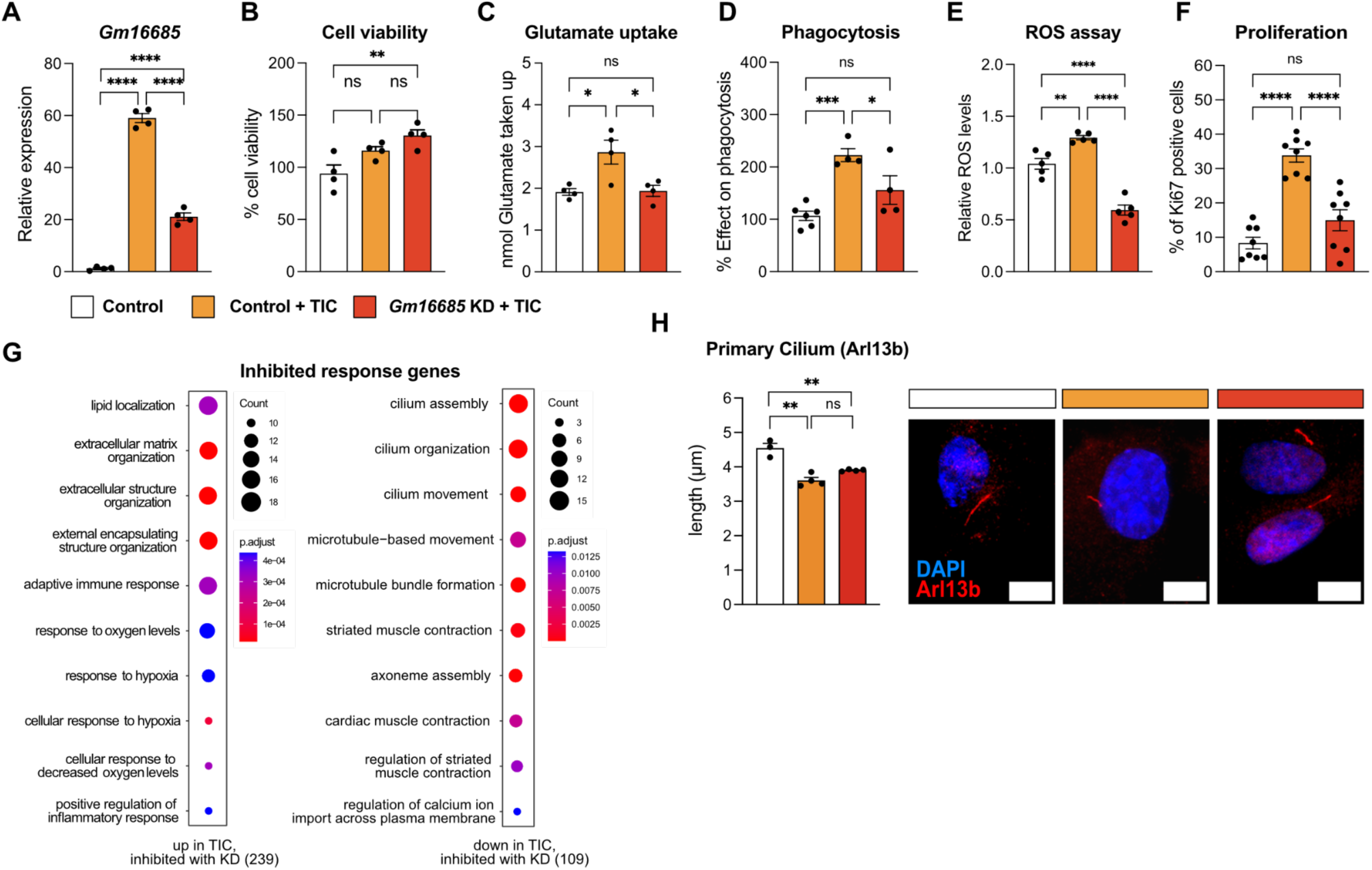
*Gm16685* knockdown inhibits reactive phenotypes in activated astrocytes. **A** Bar plot showing qPCR results for *Gm16685* after treating primary astrocytes with Gapmers to knockdown (KD) *Gm16685* in activated astrocytes. Cytokines TNF-ɑ, IL-1ɑ, C1q (TIC) or vehicle control were added 24 hours after the addition of *Gm16685* or control Gapmers and RNA was collected 24 hours later (one-way ANOVA; \*\*\*\**P* < 0.0001). Gene expression was normalized to 18S. **B-F,H** Bar plots showing results of cell viability (B), glutamate uptake assay (C), phagocytosis assay (D), reactive oxygen species (ROS) assay (E), relative amount of proliferating primary astrocytes determined by Ki67 immunofluorescence imaging (F) and length measurements of primary cilia determined by Arl13b immunofluorescence imaging (H, left panel) in primary astrocytes after *Gm16685* KD compared to TIC or untreated controls (one-way ANOVA; \**P* < 0.05, \*\**P* < 0.001, \*\*\**P* < 0.001, \*\*\*\**P* < 0.0001, ns not significant). Error bars represent SEM. **G** Gene ontology biological process analysis for TIC treatment-responsive genes showing a loss of the treatment-induced regulation upon prior *Gm16685* KD compared to control Gapmers in primary astrocytes identified from total RNA sequencing. Left panel: Genes up regulated in TIC activated astrocytes with inhibited induction upon *Gm16685* KD. Right panel: Genes down regulated in TIC activated astrocytes with inhibited reduction upon *Gm16685* KD. Analysis was done using clusterProfiler (v4.6.0). Two-sided hypergeometric test was used to calculate the importance of each term and the Benjamini-Hochberg procedure was applied for the *P* value correction. **H** Right panel: Representative immunofluorescence images of primary cilium stained with anti-Arl13b antibody. Nuclei are stained with DAPI. Scale bar 5 µm.

To delineate the underlying transcriptional changes induced by *Gm16685* KD, total RNA sequencing and differential gene expression analysis were employed (**Supplemental table 6**). We quantified transcriptional changes by comparing log2FC across two experimental contrasts: control versus TIC-treated cells, and TIC-treated versus *Gm16685* knockdown TIC-treated cells. For each gene, log2FC values were first normalized relative to the control condition to establish a common baseline. To assess the extent to which *Gm16685* knockdown inhibited gene expression toward homeostatic levels, we calculated a delta (Δ) metric defined as the difference between the TIC-induced change and the change observed upon knockdown, that captures the degree to which the knockdown counteracts TIC-driven transcriptional perturbations. Genes with a Δ value greater than 1.5 were defined as exhibiting substantial rescue toward baseline expression, indicating a strong reversal of TIC-induced effects. We found that *Gm16685* KD abolished the TIC treatment-induced upregulation (239 genes) or downregulation (109 genes), indicating that *Gm16685* is required for their treatment responsiveness (**Supplemental table 7)**. Gene ontology enrichment analysis further revealed genes with inhibited upregulation associated with ‘extracellular matrix organization’, ‘adaptive immune response’ or ‘response to decreased oxygen levels’, which is in line with our previous finding of reduced ROS production, while genes showing inhibited downregulation associated with ‘cilium assembly’, ‘cilium organization’ or ‘cilium movement’ (**Fig. 3H, Supplemental table 8**). In accordance with this, we found that the shortening of the primary cilium caused by TIC activation was less prominent in activated *Gm16685* KD astrocytes (**Fig. 3H**). In summary, our findings imply that *Gm16685* is a crucial regulator of astrocyte activation.

Next, we set out to identify the molecular mechanism of *Gm16685* during astrocyte activation, with its nuclear localization hinting towards a role in transcription. To delineate more direct transcriptional targets of *Gm16685*, we leveraged time-resolved gene expression changes and performed total RNA sequencing of 2 hour TIC activated *Gm16685* KD astrocytes with respective controls (**Supplemental table 9**). Again, we found that the knockdown of *Gm16685* abolished TIC treatment-induced upregulation (105 genes) or downregulation (106 genes) (**Fig. 4A**). When compared with the 24 hour TIC activated *Gm16685* KD astrocytes, 85 genes were still unresponsive (inhibited) while 126 genes showed delayed responsiveness (**Fig. 4A,B, Supplemental table 10**). Furthermore, transcription factor enrichment analysis from the EnrichR TRRUST transcription factor data base (https://maayanlab.cloud/Enrichr/enrich) revealed the association of these genes with STAT3 (**Fig. 4B**), which forms a signalling axis with NF-κB and helps sustain the reactive astrocyte state (Ageeva et al., 2024). In line with this, gene ontology enrichment analysis revealed the association of TIC-induced delayed responsive genes with processes like ‘response to virus’ and ‘response to interferon-beta’ (**Fig. 4C, Supplemental table 11**), while no enrichment was found for the inhibited gene subset.

**Figure 4.**
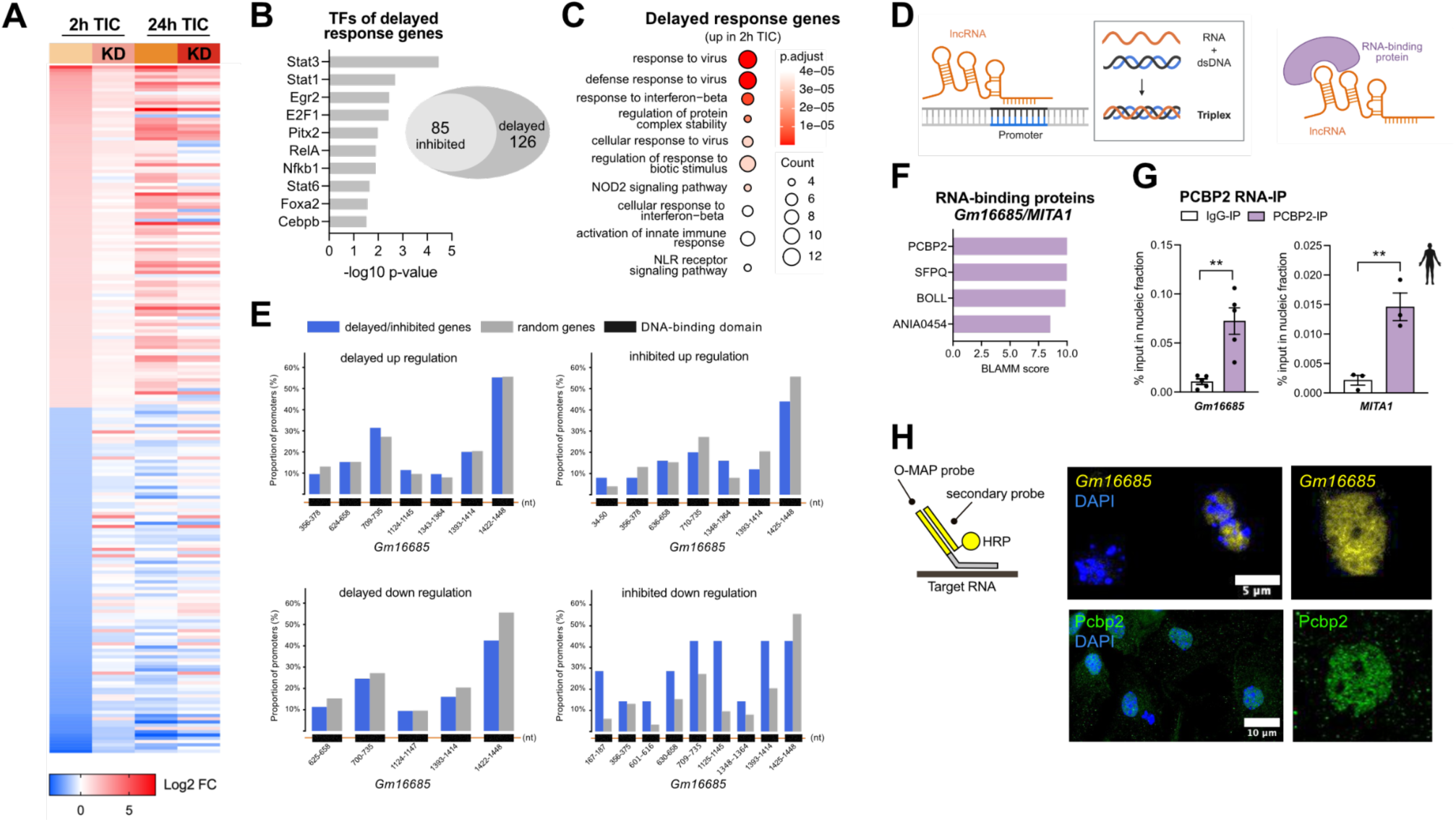
*Gm16685* knockdown delays inflammatory gene expression response in activated astrocytes. **A** Heatmap depicting Log2 fold changes (FC) of 2-hour cytokine mix TNF-ɑ, IL1-ɑ, C1q (TIC) treatment-responsive genes showing a loss of the treatment-induced regulation upon prior *Gm16685* knockdown (KD) compared to control Gapmers in primary astrocytes identified from total RNA sequencing. Values from 2h TIC, KD 2h TIC, 24h TIC and KD 24h TIC are displayed. **B** Right panel: Bar plot showing transcription factors (TF) from the EnrichR TRRUST transcription factor database (https://maayanlab.cloud/Enrichr/enrich) for delayed and inhibited genes. Left panel: Venn diagram depicting number of delayed genes (loss of treatment responsiveness at 2h TIC but not 24h TIC) or inhibited genes (loss of treatment responsiveness at 2h and 24h TIC) identified in (A). **C** Gene ontology enrichment analysis for genes upregulated in 2h TIC with delayed response in *Gm16685* KD 24h TIC. Analysis was done using clusterProfiler (v4.6.0). Two-sided hypergeometric test was used to calculate the importance of each term and the Benjamini-Hochberg procedure was applied for the *P* value correction. **D** Schematic illustration of a triplex-forming domain within a long non-coding (lncRNA, orange) engaging a homopurine stretch in double-stranded DNA (dsDNA, blue-black) via Hoogsteen base pairing in the major groove, forming an RNA-DNA:DNA triple helix (triplex) and the interaction of a lncRNA with an RNA-binding protein (violet) via structural motifs like stem loops, hairpins or single-stranded motifs. **E** Bar charts showing proportion of promoters of delayed or inhibited genes identified in (A) with a predicted DNA binding domain (DBD) for *Gm16685* from the triplex domain finder tool (Kuo et al., 2019). Statistical significance was determined using the Triplex Domain Finder promoter test with Benjamini-Hochberg false discovery rate (FDR) correction of p-values (*P < 0.05, FDR-adjusted). LncRNA sequence from 5’ to 3’ in orange, possible DNA-binding domains in black with indicated DNA nucleotide (nt) position, delayed or inhibited genes in blue, random genes in grey. **F** Bar plot showing BLAMM probability scores of RNA-binding proteins predicted to bind to both mouse *Gm16685* and human *MITA1.* **G** Bar plots depicting qPCR results from RNA immunoprecipitation of PCBP2 and IgG control for *Gm16685* in primary astrocytes (left) and *MITA1* in immortalized human astrocytes (right) treated 24h with TIC (unpaired *t* test; \*\**P* < 0.01, \*\*\*\**P* < 0.0001). Error bars represent SEM. **H** Left panel: Schematic illustration of the oligonucleotide-mediated proximity interactome mapping (O-MAP) method using a primary probe (O-MAP probe) targeting the RNA of interest and a secondary probe coupled to horseradish peroxidase (HRP). Landing pad sequence depicted in yellow. Right panel: Representative microscopy images of TIC-activated astrocytes with (top) O-MAP for *Gm16685* detected with Neutravidin, scale bar 5 µm, and (bottom) immunofluorescence staining using anti-Pcbp2 antibody, scale bar 10 µm. Nuclei were stained with DAPI.

Since lncRNAs are known to bind to DNA, RNA and proteins to exert their regulatory function (**Fig. 4D**), we first assessed if inhibited or delayed genes can potentially bind *Gm16685* via their promoter region. However, employing the Triplex Domain Finder tool (Kuo et al., 2019), we did not observe any significant DNA-binding domains in the *Gm16685* RNA (**Fig. 4E**). Next, we identified protein binding sites in the mouse *Gm16685* and human *MITA1* RNA using BLAMM (Binding site Location Analysis using Motif Models) (Fostier, 2020), which revealed four possible RNA-binding protein partners (**Fig. 4F**). Of these, two were especially of interest due to their known roles during inflammation and their localisation to the nucleus, namely splicing factor PCBP2 (Poly(rC) binding protein 2) and SFPQ (Splicing Factor Proline- and Glutamine-Rich). Employing RNA immunoprecipitation (RNA-IP), we found that *Gm16685* was enriched in Pcbp2 pulldown from nuclear lysates of activated primary astrocytes compared to IgG control, and conversely, *MITA1* was enriched in PCBP2 pulldown from nuclear lysates of activated human immortalized astrocytes (**Fig. 4G**). In line with this, oligonucleotide-mediated proximity-interactome mapping (O-MAP) imaging of *Gm16685* and immunofluorescence staining of Pcbp2 in TIC-activated primary astrocytes showed a similar nucleoplasm localisation pattern (**Fig. 4H)**.

### *Gm16685* stabilizes Pcbp2 and orchestrates Ikkβ isoform-biased target activation

Recent work showed that PCBP2 regulates splicing of *Ikbkb* mRNA in T cells, and that loss of PCBP2 increases retention of the intron between exons 8 and 9 (Martinelli et al., 2023). In our *Gm16685* KD RNA-seq data, DEXSeq analysis (Anders et al., 2012) revealed increased retention of the same intron in KD compared to control Gapmer TIC-activated astrocytes (**Fig. 5A, Supplemental table 12**). The resulting Ikkβ isoform is lacking the C-terminal helix-loop-helix (HLH) domain, which recent studies show is critical for efficient phosphorylation of IκBα and pro-inflammatory NF-κB signalling, whereas phosphorylation of AMPKα1 is retained and associated with anti-inflammatory signalling (Liu et al., 2022) (**Fig. 5A**). Consistent with this, we detected a slightly smaller Ikkβ isoform of around 90 kDa in *Gm16685* KD astrocytes that correlates with increased levels of phosphorylated Ampkα1 at Thr183 (**Fig. 5B,C**). Western blot analysis further revealed that Pcbp2 protein levels rise during TIC-induced astrocyte activation, whereas prior *Gm16685* KD reduced Pcbp2 abundance (**Fig. 5B,C**) but did not affect its mRNA levels (**Fig. 5D)**, indicating that *Gm16685* helps maintain Pcbp2 protein stability during activation. A shift of Ikkβ toward Ampkα1 phosphorylation contributes to delayed and attenuated inflammatory gene expression by reducing IκBα/NF-κB signalling while engaging an Ampk-dependent anti-inflammatory program, as seen repeatedly with *Gm16685* KD in this study. Moreover, activated AMPKα1 promotes cytoplasmic retention of the full-length STAT3 isoform (Stat3α; 90 kDa) by enhancing Ser727 phosphorylation (inhibitory) (Xiang et al., 2025) and reducing Tyr705 phosphorylation (activating) (Nerstedt et al., 2010), and the latter was recapitulated in our *Gm16685* knockdown (**Fig. 5B,C**). Moreover, we observed an increase of Stat3α upon TIC activation which decreased with prior *Gm16685* KD, while the Stat3 isoform lacking the C-terminus which acts as transcriptional repressor (Stat3β; 80 kDa) increased in *Gm16685* KD astrocytes (**Fig. 5B,C**). Consequently, Stat3-mediated pro-inflammatory gene expression is suppressed, which is consistent with Stat3 being the primary transcription factor driving the inhibited or delayed genes observed after *Gm16685* KD with 2 hours of TIC activation in **Fig. 4B**.

**Figure 5.**
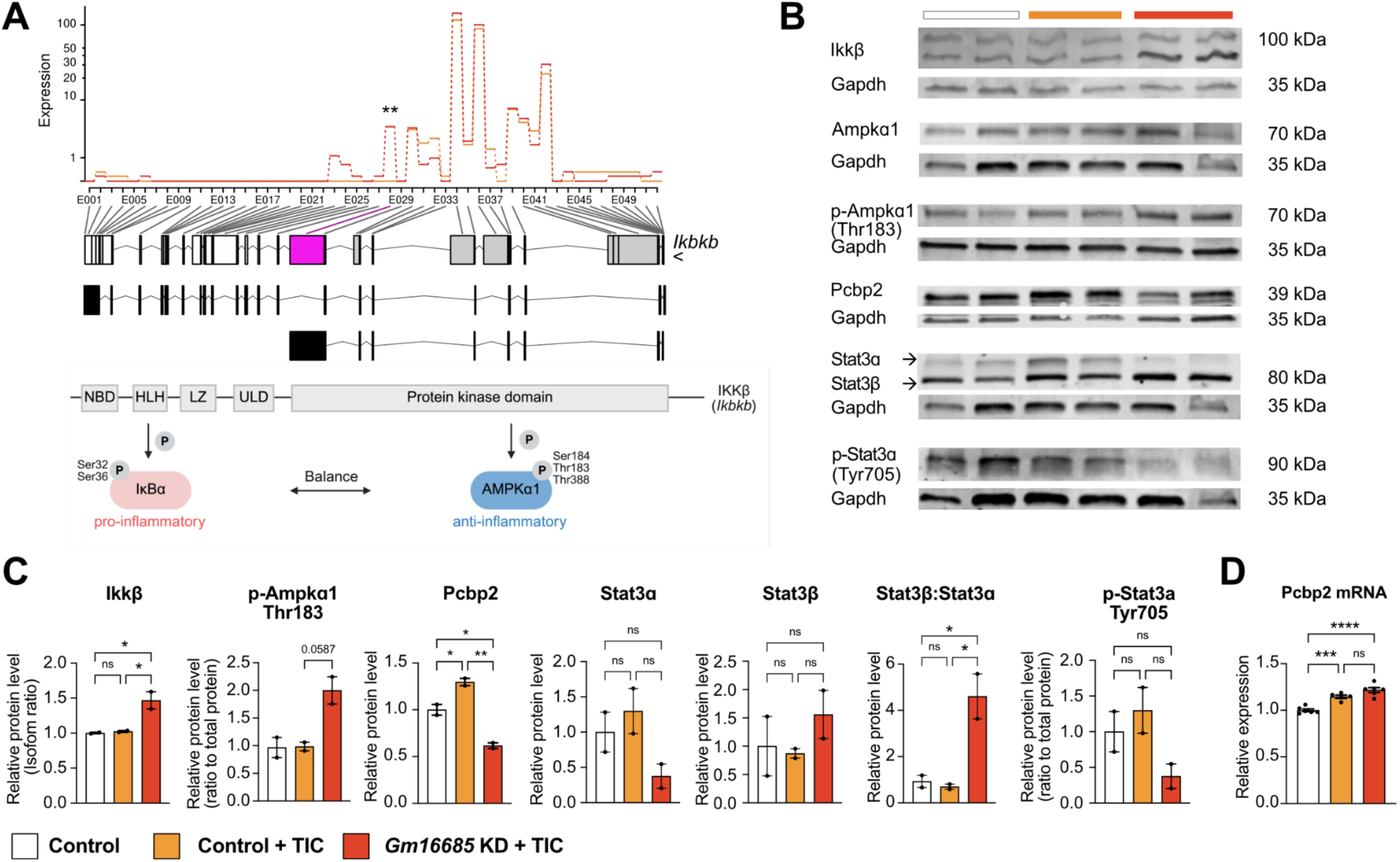
*Gm16685* stabilizes PCBP2 and orchestrates Ikkβ isoform-biased target activation. **A** Top panel: Plot showing differential exon expression for *Ikbkb* (gene coding for Ikkβ) in total RNA-seq data of 48 h Gapmer-mediated *Gm16685* knockdown (KD) in 24 h cytokine mix TNF-ɑ, IL-1ɑ, C1q (TIC) activated primary astrocytes compared to control Gapmer (Control) TIC activated astrocytes using the DEXseq package (Anders et al., 2012). Differentially expressed sites indicated in pink, resulting isoform depicted in black below. Bottom panel: Schematic illustration of functional Ikkβ domains with corresponding encoding exons indicated above and respective phosphorylation targets IkBɑ and Ampkɑ1. Nemo-binding domain (NBD); Helix-loop-helix (HLH); Leucine Zipper (LZ); Ubiquitin-like domain (ULD) (** *P* < 0.01). **B** Representative Western blots of Ikkβ, Ampkɑ1, p-Ampkɑ1 Thr183, Pcbp2, Stat3, p-Stat3ɑ Tyr705 and loading control Gapdh in whole lysates of *Gm16685* KD 24 h TIC-activated primary astrocytes compared to control Gapmers. Molecular weights (kDa) are indicated. **C** Bar plots depicting densitometric quantification of target protein levels normalized to Gapdh from (B), (one-way ANOVA or unpaired t Test; \**P* < 0.05, \*\**P* < 0.001, ns not significant). **D** Bar plot showing relative mRNA expression levels of *Pcbp2* in total RNA-seq data of *Gm16685* KD TIC-activated primary astrocytes compared to controls (one-way ANOVA; \*\*\**P* < 0.001, \*\*\*\**P* < 0.0001, ns not significant).

Taken together, our data support a model in which the lncRNA *Gm16685* stabilizes Pcbp2 protein to repress intron retention in Ikkβ pre-mRNA, thereby maintaining an Ikkβ isoform biased toward NF-κB activation (**Fig. 6**). Consequently, *Gm16685* knockdown reduces Pcbp2 levels which promotes Ikkβ intron retention, giving rise to a protein isoform which shifts its kinase activity toward Ampkα1 phosphorylation. In turn, activated Ampkα1 inhibits Stat3 activation, resulting in a delayed inflammatory response in activated astrocytes.

**Figure 6.**
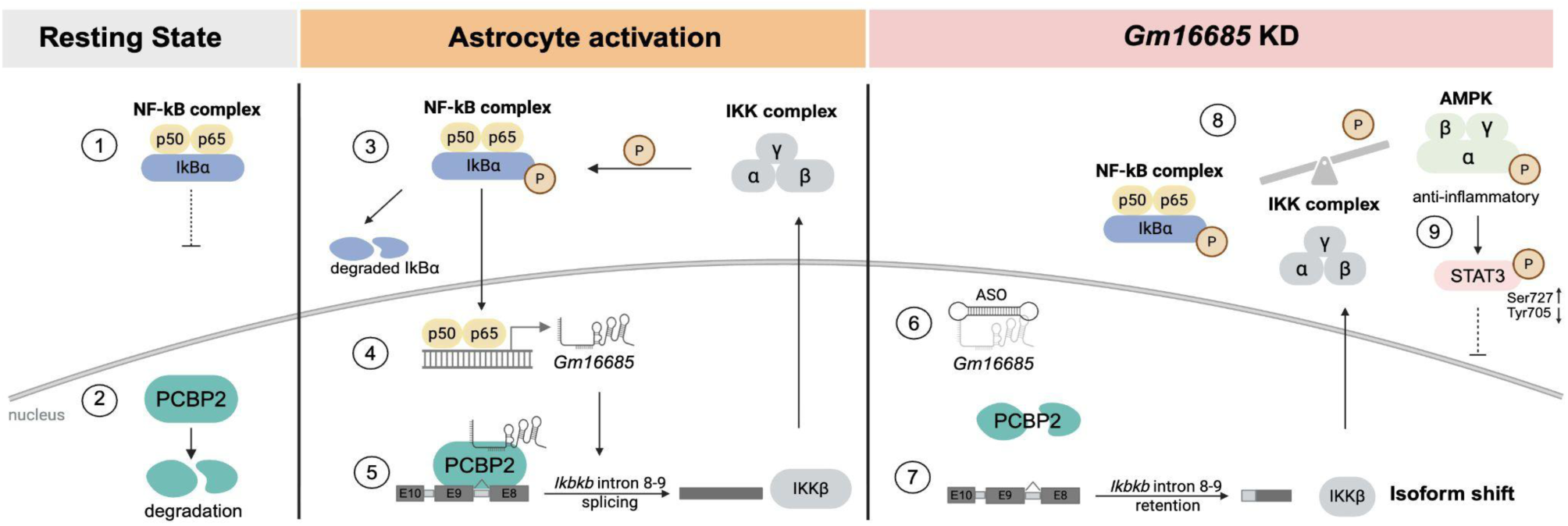
*Gm16685* mechanism of action during astrocyte activation. In the resting state, (1) inactive NF-κB (typically a p50/p65 heterodimer) is maintained in an inactive state in the cytoplasm by binding to the inhibitory protein IkBɑ. (2) RNA-binding protein Poly(rC)-binding protein 2 (Pcbp2) is degraded in the nucleus. Upon astrocyte activation (3) IkBɑ gets phosphorylated by the IkB kinase (IKK) complex, resulting in its degradation and active NF-κB can translocate to the nucleus to activate its target genes such as (4) the lncRNA *Gm16685.* (5) *Gm16685* binds to and stabilizes Pcbp2. Pcbp2 splices the intron between exon 8 and exon 9 in the *Ikbkb* pre-mRNA, resulting in the full-length Ikkβ isoform, responsible for IkBɑ phosphorylation and pro-inflammatory signalling. (6) Gapmer antisense-oligonucleotide (ASO)-mediated *Gm16685* knockdown (KD) prior to astrocyte activation reduces Pcbp2 protein levels in the nucleus, (7) causing *Ikbkb* intron retention resulting in a shorter Ikkβ isoform that is (8) biased towards Ampkɑ1 phosphorylation and anti-inflammatory signalling. Here, activated AMPKα1 promotes cytoplasmic retention of STAT3 by enhancing Ser727 phosphorylation (inhibitory) and reducing Tyr705 phosphorylation (activating). Created with Biorender.

## Supplementary Figures

**Figure S1.**
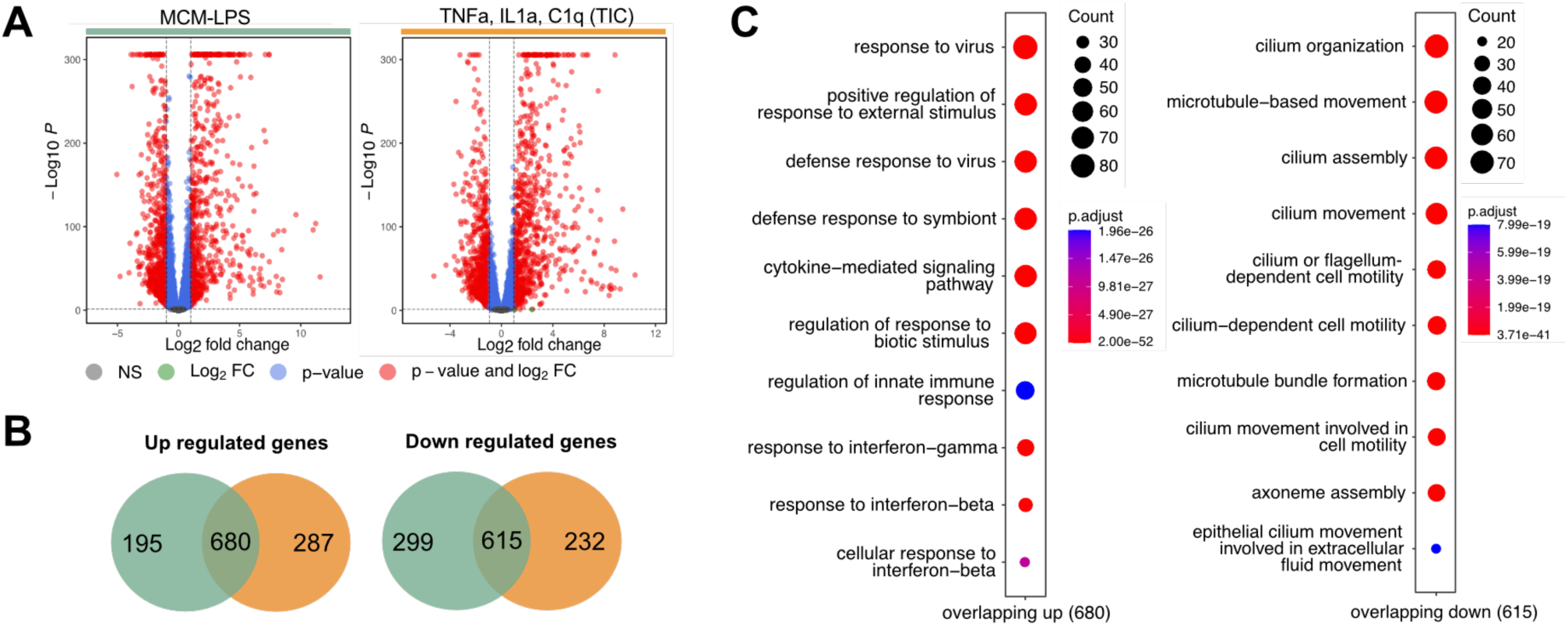
Transcriptomic comparison of astrocyte activation by microglia and cytokines. **A** Volcano plots show differentially expressed genes from total RNA sequencing of mouse primary astrocytes treated 24h with MCM-LPS vs. vehicle (left panel) or the cytokine mix TNF-ɑ, IL-1ɑ, C1q (TIC) vs. vehicle (right panel). Significant genes (adjusted p-value < 0.05 and |log2FC| > 1) are highlighted in red. **B** Venn diagrams showing overlap between MCM-LPS or cytokine mix differentially expressed genes that are significantly up regulated (left panel) or significantly down regulated (right panel), (adjusted p-value < 0.05, |log2FC| > 1 and BaseMean > 25)**. C** Plots showing the results of a GO term analysis for selected genes displayed in (A) and (B). Left panel: up regulated overlapping genes. Right panel: down regulated overlapping genes. Analysis was done using clusterProfiler (v4.6.0). Two-sided hypergeometric test was used to calculate the importance of each term and the Benjamini-Hochberg procedure was applied for the *P* value correction.

**Figure S2.**
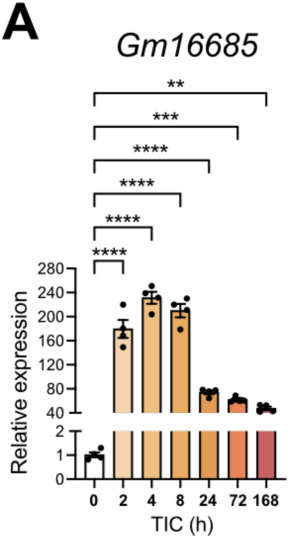
**A** Bar plot showing qPCR results for *Gm16685* in primary astrocytes treated with the cytokine mix TNF-ɑ, IL-1ɑ, C1q (TIC) ranging from 2 hours (h) until 7 days (168 h) compared to vehicle control (0 h). Gene expression was normalized to 18S (one-way ANOVA, \*\**P* < 0.01, \*\*\**P* < 0.001, \*\*\*\**P* < 0.0001). Error bars represent SEM.

## Discussion

In this study, we describe a layer of deregulated lncRNAs in a classic microglia-driven paradigm of astrocyte activation (Liddelow et al., 2017). We show that a subset of evolutionarily conserved lncRNAs is already induced after 2 hours of activation and that brain-specific lncRNAs are prone to decrease in expression. We found that the lncRNA *Gm16685* is strongly induced in activated astrocytes and that its loss attenuates inflammatory phenotypes and gene expression, with mechanistic evidence linking this effect to PCBP2, *Ikbkb* mRNA processing and altered downstream inflammatory signalling.

Astrocyte reactivity plays a central, disease-specific role across neurodegenerative disorders including AD, PD, and FTD (Escartin et al., 2021) (Li et al., 2019). Here, reactive astrocytes act as responders to proteinopathy and are active contributors to neuroinflammation and synaptic dysfunction. Consistent with potential disease relevance, *MITA1* expression was increased in selected human datasets from AD, PD and FTD. While physiological activation supports repair, pathological states driven by amyloid beta/tau in AD, α-synuclein in PD, or TDP-43/progranulin deficits in FTD, engage shared signalling hubs like JAK/STAT3 and NF-κB pathways (Ben Haim et al., 2015), (Li et al., 2019), resulting in the increased secretion of pro-inflammatory cytokines (Escartin et al., 2021). Preclinical trials have started to approach targeting astrocyte reactivity, especially NF-κB or Stat3 signalling pathways, in AD mouse models (Reichenbach et al., 2019), (Nakano-Kobayashi et al., 2023), (Kong et al., 2014), positioning these regulators as promising therapeutic targets across proteinopathies. However, most studies target the protein coding genome.

In the last decade, lncRNAs have emerged as critical regulators of cellular mechanisms, especially in the CNS. Thus, understanding the intricate interplay between the non-coding and coding genome is crucial to our understanding of disease-associated astrocytes in the context of neurodegeneration. Our study shows that lncRNAs downregulated in activated astrocytes display higher brain-specificity compared to upregulated lncRNAs, indicating a shift from lncRNAs involved in homeostasis and cellular identity to lncRNAs with broader expression patterns and immune functions in disease. Notably, several of the identified lncRNAs have been previously implicated in inflammatory or glial responses, including *Prdm16os* in astrocyte function and reactive states (Schröder et al., 2024), *AW112010* in CNS and systemic inflammatory signalling (Yang et al., 2020), *Meg3* in microglial Nlrp3-mediated inflammation (Meng et al., 2021), and *Rmst* in neuron-glia associated inflammatory and neuropathic pain pathways (Zhang et al., 2025). Our work further uncovered 14 evolutionary conserved lncRNAs that show expression changes already in the initial phase of 2-hour astrocyte activation.

Here, we focus on *Gm16685*, a low abundant lncRNA in resting astrocytes that is strongly upregulated upon astrocyte activation. Indeed, work on *Gm16685* has previously been published in the context of other inflammation-induced diseases like colitis (Akıncılar et al., 2021), liver *S. japonicum* infection (Zhao et al., 2023) or HIV-associated neurocognitive disorder (Zhou et al., 2018), while its human homolog, *MITA1* (metabolism-induced tumor activator), was first described induced by energy stress in hepatocellular carcinoma (Ma et al., 2019). Taken together, this supports a crucial role of *Gm16685/MITA1* in the inflammatory response of different tissues.

Furthermore, we hypothesised that perturbations in the identified lncRNA subset during the initial phase of astrocyte activation would cause shifts in reactive phenotypes. Indeed, knockdown of *Gm16685* caused the abolishment of different reactive phenotypes like increased glutamate uptake, elevated phagocytosis or increased ROS levels during astrocyte activation. Acute phenotypes like enhanced phagocytosis are primarily considered beneficial, leading to decreased amounts of cell debris or protein aggregates (Jiwaji et al., 2022), while an increase in ROS is considered detrimental, contributing to the dysfunction of activated astrocytes (Sheng et al., 2013). The interpretation of altered glutamate uptake is more complex, because enhanced astrocytic glutamate uptake may reflect both a reactive state and a compensatory protective mechanism to limit extracellular glutamate accumulation.

In order to investigate the underlying molecular mechanisms of *Gm16685* in astrocytes, we employed time-resolved knockdown strategies to reduce secondary transcriptional effects (Much et al., 2024). We observed that *Gm16685* KD delayed the induction of Stat3-target genes involved in the innate immune response. Moreover, we used BLAMM to identify protein-binding motifs in our target lncRNA and found that RNA-binding protein Pcbp2 was indeed binding to both mouse *Gm16685* and human *MITA1* RNA in nuclear lysates from activated astrocytes. Intriguingly, we observed that Pcbp2 protein levels increased in TIC-activated astrocytes, while protein levels decreased upon knockdown of *Gm16685.* This finding is in line with a previous publication that discovered elevated PCBP2 levels in AD patients (Zhang et al., 2022). Furthermore, reducing PCBP2 levels mitigated AD pathology and cognitive decline in a study by Wang et al. (Wang et al., 2025). In addition, Pcbp2 deficiency decreased proliferation in spinal cord injury astrocytes (Mao et al., 2016). In parallel, *Gm16685* knockdown was associated with altered *Ikbkb* mRNA processing, including retention of the intron between exons 8 and 9, a splicing event previously linked to PCBP2 in T cells (Martinelli et al., 2023). Previous work has shown that IKKβ can signal through distinct substrate interactions, with the kinase domain mediating AMPKα1 phosphorylation and the helix loop helix region supporting IκBα phosphorylation and canonical NF-κB activation (Liu et al., 2022). Our data are consistent with a model in which *Gm16685* knockdown promotes an IKKβ isoform shift that favours AMPKα1 associated signalling over IκBα/NF-κB signalling, thereby contributing to attenuated STAT3 associated inflammatory gene expression.

A key limitation of this study is our reliance on primary cell cultures rather than *in vivo* mouse models or more complex human 3D organoids, which limits insights into multicellular interactions and tissue-level pathology. Moreover, human astrocyte models like the immortalized human astrocyte cell line used in this study often lack key functional properties and only cover some functional properties of astrocytes (Galland et al., 2019). Additionally, the TIC cytokine mix employed here only partially mimics astrocyte activation, whereas disease-associated states (e.g., in AD, PD, ALS/FTD) exhibit greater divergence (Escartin et al., 2021), (Santiago-Balmaseda et al., 2024), (İlgün & Çakır, 2025), (Gong et al., 2025), (Trias et al., 2018). This is compounded by astrocyte heterogeneity across subtypes observed in mouse and human (Batiuk et al., 2020), (Hasel et al., 2021). Single-cell RNA (scRNA) sequencing of activated astrocytes could clarify lncRNA distribution and networks within specific subpopulations. However, scRNA or snRNA sequencing data are currently not optimised to recapitulate lncRNA expression. Technical limitations of snRNA sequencing include low capture efficiency for lowly expressed, nuclear-depleted lncRNAs, incomplete annotation of brain-specific transcripts and variability in postmortem tissue quality, brain region sampling, or disease stage representation. Hence, simplified *in vitro* cell culture models and disease-mimicking perturbations enable a first glimpse into the non-coding regulatory layer involved in complex neurodegenerative diseases.

## Methods

### Animals

All animal experiments were approved by the local Animal Welfare Office of Goettingen University and the Lower Saxony State Office for Consumer Protection and Food Safety. Pregnant CD-1 mice were obtained from Janvier Labs. Animals were housed in standard cages with a 12-h dark and light cycle. Water and food were provided ad libitum.

### Primary astrocyte culture

Primary astrocyte cultures were generated from postnatal day 1 mouse cortices and hippocampi as previously described (Schröder et al.). Briefly, tissue was dissociated with 0.05 % trypsin-EDTA, and cells from 2-3 pups were plated on poly-D-lysine-coated flasks in fetal bovine serum-containing DMEM, then shaken 7 days later to remove non-astrocytic cells and re-plated in Neurobasal Plus medium supplemented with B-27 Plus (Gibco), GlutaMAX (Gibco), penicillin/streptomycin (Gibco), and heparin-binding EGF-like growth factor (Sigma-Aldrich) at 15,000 cells/cm². Cultures were maintained at 37°C, 5% CO₂ with half-medium changes twice per week.

### Immortalized human fetal astrocytes

Immortalized human fetal astrocytes (hTERT; abm) were obtained at a density of >1 × 10⁶ cells/ml and expanded in culture according to the supplier’s recommendations. Experiments were conducted at 80% confluency.

### Stimulation of astrocytes

Primary microglia were cultured as previously described (Islam et al., 2021) and stimulated as previously shown (Schröder et al., 2024). In brief, microglia were activated with 100 ng/mL LPS for 4 hours. The medium was then renewed and collected by filtering after 6 hours. On DIV13, primary astrocytes were treated with a 1:1 mixture of microglial conditioned medium and fresh medium, and RNA was isolated 24 hours later.

To mimic the activation of astrocytes by microglia (Liddelow et al., 2017), astrocytes were treated with a cytokine mixture of IL-1α (3 ng ml^−1^, Sigma, I3901), TNFα (30 ng ml^−1^, Cell Signaling Technology, 8902SF) and C1q (400 ng ml^−1^, MyBioSource, MBS143105) for 24 hours.

### Antisense LNA Gapmers

Antisense LNA Gapmers targeting *Gm16685, MITA1* or negative controls (NC) were designed and purchased from Qiagen having the following sequences:

NC: GCTCCCTTCAATCCAA

*Gm16685 (1)*: GGTCGGTTAAAGTTAG

*Gm16685 (2)*: CCAGCACGTCATTCTA

*MITA1*: TTAGAGAGGTTGGAGT

Transfection was performed using Lipofectamine RNAiMAX (Thermo Fisher Scientific) according to the manufacturer’s instructions. Cells were treated on DIV12 and all experiments were performed on DIV14. Cell viability was measured using PrestoBlue (Thermo Fisher Scientific).

### Glutamate uptake

On DIV14, primary astrocytes were incubated with 100 µM glutamate for 1 hour as previously described (Schröder et al., 2024). Supernatant was collected and the amount of remaining glutamate was determined using the Glutamate-Glo™ Assay (Promega) following the manufacturer’s instructions. Luminescence was recorded with a FLUOstar® Omega (BMG).

### Lactate secretion

To estimate lactate secretion, the complete medium was changed 24 hours before the endpoint of an experiment. The medium was collected and lactate concentrations were measured using the Lactate-Glo™ Assay (Promega) following the manufacturer’s instructions. Luminescence was recorded with a FLUOstar® Omega (BMG).

### Phagocytosis

Phagocytic activity of primary astrocytes was quantified using the Vybrant Phagocytosis Assay (Invitrogen). Astrocytes were seeded on black-walled 96-well glass-bottom plates at 7,000 cells per well, and empty wells containing only culture medium served as no-cell controls. After the respective treatments, a fluorescent E. coli BioParticle suspension was prepared according to the manufacturer’s instructions, added to each well, and cells were incubated for 2 h at 37°C. The BioParticle suspension was then removed and extracellular fluorescence quenched by a 1 min incubation with trypan blue solution, followed by two PBS washes. Intracellular fluorescence was measured at 480 nm excitation/520 nm emission on a FLUOStar Omega plate reader (BMG). The percentage effect of a treatment on phagocytic activity was calculated by subtracting the no-cell control signal from the treatment signal and dividing this value by the difference between the cell control signal and the no-cell control signal.

### Reactive oxygen species

Intracellular reactive oxygen species (ROS) were quantified using the DCFDA/H2DCFDA Cellular ROS Assay Kit (Abcam), according to the manufacturer’s protocol. Primary astrocytes were seeded on black, clear-bottom 96-well plates at 4,000 cells per well on DIV7 and subjected to the indicated treatments from DIV12. On DIV14, ROS activity was determined with fluorescence signal measurement, recorded with a FLUOstar® Omega microplate reader (BMG).

### Immunofluorescence

Immunofluorescence staining was used to assess both cell proliferation (Ki67) and primary cilia length (Arl13b) in cultured astrocytes. Cells were grown on chamber slides, fixed in 4% paraformaldehyde for 10 min at room temperature and washed once with PBS. For Ki67 staining, cells were permeabilized with 0.1% Triton X-100 in PBS (PBS-T) for 15 min, then blocked for 1 h in PBS containing 2% BSA, 20% normal goat serum (NGS), and 0.1% Tween-20, followed by overnight incubation at 4°C with anti-Ki67 (1:1000; Abcam) diluted in 2% BSA and 5% NGS in PBS-T and 1 h incubation with fluorophore-conjugated secondary antibody in 2% BSA and 1.5% NGS in PBS-T. For Arl13b staining, slides were blocked for 1 h at room temperature in PBS containing 10% goat serum and 0.1% Triton X-100, followed by overnight incubation at 4°C with anti-Arl13b (1:1000; Proteintech) in the same solution and 1 h incubation with fluorophore-conjugated secondary antibody (1:1000). Finally, slides were incubated with DAPI for 5 min, and mounted with ProLong Gold antifade reagent. Images were acquired on a Keyence BZ-X800 microscope using 20× or 40× objectives.

### Cytoplasmic and nuclear fractionation

Cytoplasmic and nuclear fractions were prepared as previously described (Schröder et al., 2024). In brief, DIV14 primary astrocytes were fractionated using the EZ prep lysis kit (Sigma-Aldrich) according to the manufacturer’s protocol, supplemented with RNase inhibitor (Promega), followed by TRIzol LS (Thermo Fisher Scientific) extraction and storage at -20°C.

### Oligonucleotide-mediated proximity interactome mapping imaging

Oligonucleotide-mediated proximity interactome mapping (O-MAP) imaging was performed essentially as described before (Tsue et al., 2024). Primary astrocytes were seeded onto poly-D-lysine-coated glass coverslips and cultured to DIV14 before fixation with 2% formaldehyde, permeabilization with Triton X-100, and inactivation of endogenous peroxidases. *Gm16685* RNA was detected using 1 µM custom oligonucleotide pools (oPools; IDT) (**Supplementary table 13**) and hybridized overnight in a 10 % formamide-containing buffer at 37°C. For imaging experiments, endogenous biotinylated proteins were blocked and free streptavidin binding sites were saturated with d-biotin (Thermo Fisher Scientific). Samples were incubated with SABER1-HRP secondary probe (100 nM; Bio-Synthesis) in 30 % formamide-containing buffer for 1 h at 37°C, and subjected to 10 min *in situ* biotinylation with labelling solution containing biotinyl-tyramide (Sigma Aldrich) and H₂O₂, followed by quenching, SABER2-AF647 (IDT) RNA-FISH, NeutrAvidin Protein DyLight™ 550 (Thermo Fisher Scientific) staining, DAPI counterstaining, and mounting in ProLong Gold antifade mountant (Invitrogen). Images were acquired on a Keyence BZ-X800 microscope (Keyence, Japan) using a 60x oil objective and analysed in Fiji/ImageJ.

### RNA immunoprecipitation

RNA immunoprecipitation (RNA-IP) was performed essentially as described recently (Schröder et al., 2024). In brief, primary astrocytes were cultured in T75 flasks and, at DIV14, dissociated with 0.25% trypsin-EDTA, washed in PBS, and fractionated into cytoplasmic and nuclear lysates using NP-40-based hypotonic buffer followed by sucrose-containing TSE buffer with protease and RNase inhibitors. Nuclear pellets were sonicated, clarified by centrifugation, and lysates were quantified by BCA assay and pre-cleared with Protein A/G magnetic beads. First, 10 % of input was taken, then PCBP2-RNA complexes were immunoprecipitated overnight at 4°C using 3 µg of anti-PCBP2 antibody (PA5-22350, Invitrogen) or IgG isotype control (Cell Signaling Technologies) bound to Protein A/G beads in RNA-IP buffer, washed, and eluted by proteinase K treatment. RNA was then purified using the RNA clean & concentrator-5 kit (Zymo Research) according to the manufacturer’s instructions.

### RNA extraction

Total RNA was extracted from cells using TRI Reagent (Sigma-Aldrich) and the RNA Clean & Concentrator-5 kit (Zymo Research) per manufacturer’s instructions. Concentrations were quantified by NanoDrop or Qubit (RNA HS Assay; Thermo Fisher Scientific), and quality assessed by Bioanalyzer (Agilent) for sequencing samples.

### cDNA, qPCR

cDNA was synthesized from 20–800 ng RNA using the Transcriptor First Strand Kit (Roche) with random hexamers, diluted 10-fold, and analysed by qPCR with LightCycler 480 SYBR Master Mix (Roche) in duplicates on a LightCycler 480. Expression of non-coding and protein-coding genes was normalized to 18S (primers listed in **Supplementary Table 14**), using the 2^−ΔΔCt method (Livak & Schmittgen, 2001).

### Library preparation and total RNA sequencing

Libraries were prepared from 300 ng total RNA using the SMARTer Stranded Total RNA Sample Prep Kit-HI Mammalian (Takara Bio) following the manufacturer’s instructions, amplified for 13 cycles, and sequenced on a NextSeq 2000 (Illumina; single-end).

### Bioinformatic analysis

Raw reads were processed with bcl2fastq (v2.20.2), quality-checked with FastQC (v0.11.5), aligned to mm10 using STAR (v2.5.2b), and quantified with featureCounts (v1.5.1). Differential expression analysis was performed using DESeq2 (v1.38.3) (Love et al., 2014) with RUVSeq (v1.32.0) correction (Risso et al., 2014), followed by GO analysis with clusterProfiler (v4.6.0) (Yu et al., 2012) and transcription factor enrichment via Enrichr (https://maayanlab.cloud/Enrichr/). Differential exon usage and alternative splicing was analysed using DEXseq (v1.48.0) (Anders et al., 2012).

Computational analyses were performed to identify RBP binding sites within the mouse lncRNA *Gm16685* (ENSMUST00000348221.1) and its human homolog *MITA1* (ENST00000649603.2). Binding sites were identified using BLAMM (Binding site Location Analysis using Motif Models) (Fostier, 2020). For each lncRNA transcript, BLAMM utilized Position Weight Matrices (PWMs) to calculate a log-likelihood score, representing the strength of the match between the transcript sequence and a canonical RBP binding motif. The BLAMM score reflects the statistical match between the RNA sequence and the RBPs Position Weight Matrix (PWM). Higher scores indicate a stronger alignment with the canonical binding motif; positive scores generally represent a high likelihood of a sequence-specific interaction, while lower or negative scores suggest a weak or non-specific match, with intermediate values indicating moderate binding potential.

### Statistical analysis

Statistical analysis was performed in GraphPad Prism 9, with data presented as mean + SEM unless otherwise indicated. Two-tailed unpaired t-tests or one-way ANOVA with Tukey’s post-hoc were used as appropriate. GO and pathway enrichments employed Fisher’s exact test with Benjamini-Hochberg correction.

## Acknowledgements

AF was supported by the DFG (Deutsche Forschungsgemeinschaft) priority program 2502 EPIADAPT, SFB1286. The German Federal Ministry of Research, Technology and Space (BMFTR) via the ERA-NET Neuron project EPINEURODEVO; The EU Joint Programme Neurodegenerative Diseases (JPND)-EPI-3E; Germany’s Excellence Strategy-EXC 2067/1 390729940. FS was supported by the GoBIO project miRassay (16LW0055) by the BMFTR. UF was supported by the Hans und Ilse Breuer Stiftung and the International Max-Planck Research School (IMPRS) Genome Science Göttingen.

## Author contributions

UF planned and conducted the majority of the experiments and wrote the manuscript. TP and DMK helped with statistical analysis. SS, FS, SB and ALS coordinated and performed RNA-seq experiments. AF conceived the project, supervised progress, and wrote the manuscript.

## Data availability

RNA-sequencing will be available via the GEO database for murine data.

